# Pseudofinder: detection of pseudogenes in prokaryotic genomes

**DOI:** 10.1101/2021.10.07.463580

**Authors:** Mitchell J. Syberg-Olsen, Arkadiy I. Garber, Patrick J. Keeling, John P. McCutcheon, Filip Husnik

## Abstract

Prokaryotic genomes are generally gene dense and encode relatively few pseudogenes, or nonfunctional/inactivated remnants of genes. However, in certain contexts, such as recent ecological shifts or extreme population bottlenecks (such as those experienced by symbionts and pathogens), pseudogenes can quickly accumulate and form a substantial fraction of the genome. Identification of pseudogenes is, thus, a critical step for understanding the evolutionary forces acting upon, and the functional potential encoded within, prokaryotic genomes. Here, we present Pseudofinder, an open-source software dedicated to pseudogene identification and analysis. With Pseudofinder’s multi-pronged, reference-based approach, we demonstrate its capacity to detect a wide variety of pseudogenes, including those that are highly degraded and typically missed by gene-calling pipelines, as well newly formed pseudogenes, which can have only one or a few inactivating mutations. Additionally, Pseudofinder can detect intact genes undergoing relaxed selection, which may indicate incipient pseudogene formation. Implementation of Pseudofinder in annotation pipelines will not only clarify the functional potential of sequenced microbes, but will also generate novel insights and hypotheses regarding the evolutionary dynamics of bacterial and archaeal genomes.

## Background

Pseudogenes are remnants of genes that have fixed inactivating nucleotide substitutions or insertions/deletions relative to their ancestral coding sequences (Ochman and Davalos, 2006; Lerat and Ochman, 2005). In eukaryotic genomes, pseudogenes frequently arise from relaxed selection on one copy of a gene resulting from gene (or whole genome) duplications, and much effort has gone towards specific studies, tools, and databases to identify them (Karro *et al*., 2007; Pink *et al*., 2011). In contrast, genomes of Bacteria and Archaea are usually gene dense and encode very few pseudogenes (Kuo *et al*., 2009, Kuo and Ochman 2010, Goodhead and Darby, 2015). However, pseudogenes do exist in prokaryotic genomes (Liu et *al*., 2004; Lerat and Ochman, 2005), most commonly in species where large numbers of genes have become unnecessary through rapid and sustained changes in ecological context (Ochman and Davalos, 2006). Classic examples include intracellular bacterial endosymbionts or pathogens, where, in extreme cases, pseudogenes can outnumber functional genes (Toh et *al*., 2006; Singh and Cole, 2011; Clayton et *al*., 2012; Burke and Moran, 2011; McCutcheon and Moran, 2012; Oakeson et al., 2014).

Identification of pseudogenes is critical for understanding the physiology, metabolism, and evolutionary adaptations of pathogens and symbionts, as well as being an underappreciated step in the annotation of even “normal” bacterial genomes. For example, pseudogene annotation is important for bacterial and archaeal phylogenomics; including pseudogenes in phylogenetic trees may lead to artifacts, like overestimated branch lengths. Despite the importance of pseudogene identification, it is still commonplace for pseudogenes to be annotated manually based on arbitrary criteria, by the use of custom unpublished scripts, or by relying on automatic annotation tools, such as the NCBI (National Center for Biotechnology Information) prokaryotic genome annotation pipeline (PGAP, Tatusova et al., 2016) or DFAST (Tanizawa et al., 2018). These tools, designed primarily for functional annotation, are less-than-ideal for pseudogene prediction because they lack standardization and do not allow for species-specific adjustments.

Here we present Pseudofinder, an open-source and highly customizable program that differentiates candidate pseudogenes from intact genes. Pseudogene identification is guided by a reference-based approach where a genome-of-interest is annotated by comparison to a user-supplied protein sequence database (e.g. RefSeq [Pruitt et al., 2007]) and/or a closely related reference genome. Using the reference database of proteins, Pseudofinder makes evidence-based annotations of truncated, fragmented, and highly degraded genes. When a reference genome of suitable evolutionary distance is available, Pseudofinder has the capacity to detect cryptic pseudogenes, and reports on the type and quantity of inactivating mutations (e.g. nonsense mutations, frameshift-inducing indels, etc.).

## Results and Discussion

We tested Pseudofinder with two bacterial genomes: 1) *Ca*. Sodalis pierantonius str. SOPE (Oakeson et al., 2014, hereafter SOPE), a host-beneficial intracellular symbiont known to have many pseudogenes, and 2) *Shewanella* sp. ZOR0012 (Lebov et al., 2020, hereafter, ZOR0012), a strain closely related to S. *oneidensis* MR-1, known to inhabit zebra fish intestinal tracts. ZOR0012 is not expected to encode many pseudogenes in its genome, but as it moved to an ecological niche that is not commonly occupied by this genus, it may be undergoing different selective pressures relative to other metal-reducing *Shewanella* spp., particularly *Shewanella* MR-1, its closest known relative.

We compared pseudogene predictions from Pseudofinder to those derived from two annotation pipelines that include pseudogene prediction as part of their workflow: PGAP (Tatusova et al., 2016) and DFAST (Tanizawa et al., 2018). Because DFAST provides the option of annotation using two different geneprediction software packages (Prodigal [Hyatt *et al*., 2010] and MetaGeneAnnotator [Noguchi *et al*., 2008]), which may differ in genes predicted, we ran DFAST using both gene-calling methods. Additionally, it is worth noting that PGAP uses GeneMark for gene prediction, which may also result in differences in gene predictions. In light of these potential differences, we included in our benchmarking only those genes that were predicted by all three gene-calling pipelines. This means that we excluded pseudogene candidates identified in intergenic regions (i.e. regions between genes where no open reading frame is detected). Pseudofinder analysis was carried out using *Shewanella* MR-1 and *Sodalis praecaptivus* HS as reference genomes for ZOR0012 and SOPE, respectively.

In both the SOPE and ZOR0012 genomes, Pseudofinder predicted the greatest number of pseudogenes compared to PGAP and DFAST. Of all Pseudofinder-identified pseudogenes, 87.6% (ZOR0012) and 65.8% (SOPE) were also flagged by at least one other annotation software (**Figure 1A**). The remaining 12.4% and 34.2%, which can be considered Pseudofinder-specific pseudogene candidates, were flagged by Pseudofinder for a range of different reasons, the most common being elevated *dN/dS* (**Figure 1B**), a metric that is not used by other annotation pipelines. SOPE, specifically, encodes a large number of genes experiencing relaxed selection (Oakeson *et al*., 2014), which make up 625 of the 703 Pseudofinder-specific pseudogene candidates in that genome. Genes considerably shorter than their top homologs from the reference database were also relatively common among the Pseudofinder-specific pseudogenes. Additionally, Pseudofinder identified gene remnants in genomic regions where no open reading frame was predicted (i.e. intergenic regions): 238 in ZOR0012 and 305 in SOPE, but these counts are not included in the numbers presented in **Figure 1**.

**Figure 1:**
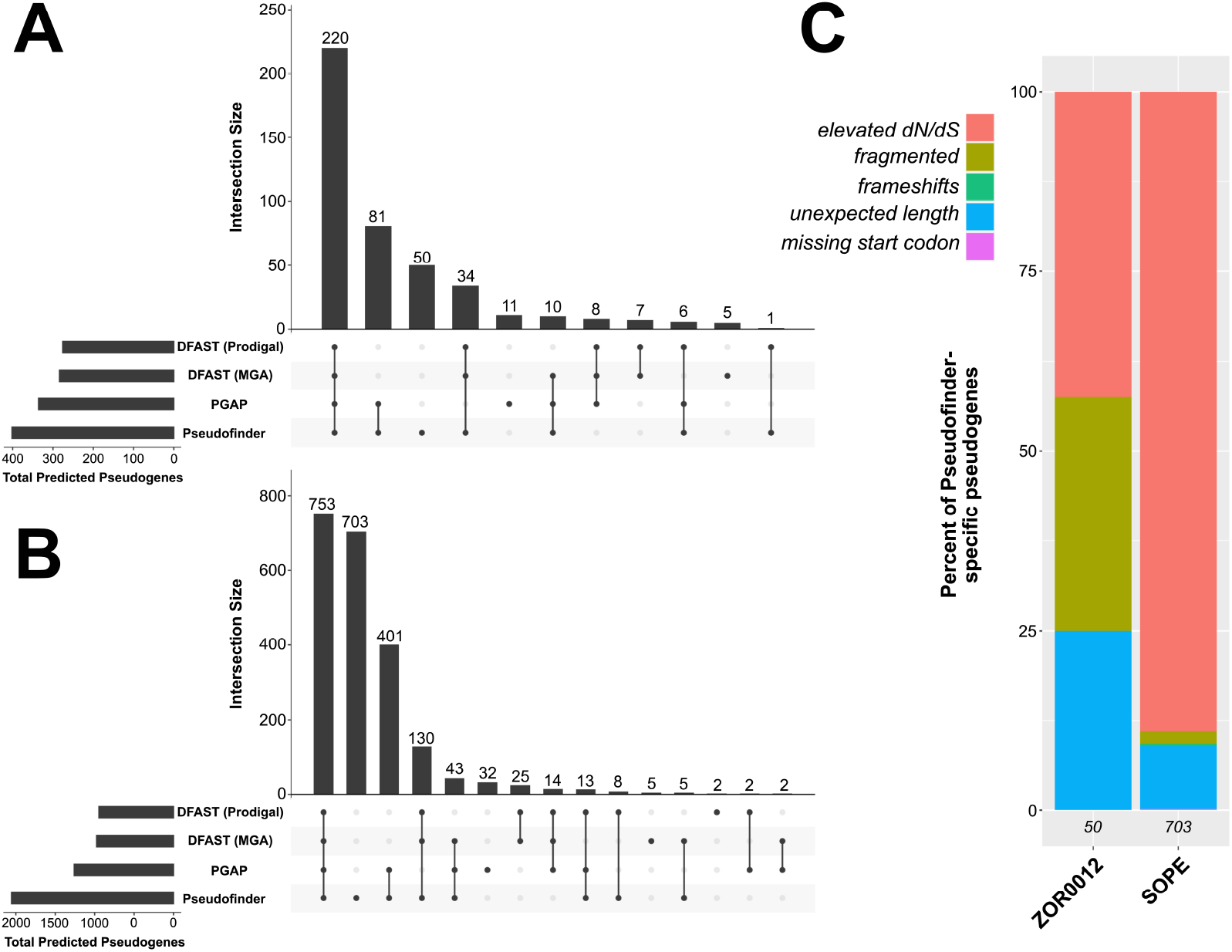
Summary of benchmarking results, comparing pseudogene predictions by Pseudofinder to those of two other software: PGAP and DFAST (run with two different gene-callers). A and B) ‘Upset’ plots (Conway et al., 2017), showing the overlap and differences between the three pipelines in pseudogenes predicted from ZOR0012 (A) and SOPE (B). Each bar in the barplot represents the total number of pseudogenes that overlap between the pipelines denoted with dots below. C) Barplot showing the types of pseudogenes that were predicted only by Pseudofinder in SOPE and ZOR0012 (i.e. Pseudofinder-specific pseudogenes). Italicized numbers at the bottom of each bar indicate the number of Pseudofinder-specific pseudogenes included in each genome.

Genes predicted to be pseudogenes by either or both PGAP or DFAST, but missed by Pseudofinder, represent 7.2% (ZOR0012) and 3.8% (SOPE) of the total predicted pseudogenes from each genome. PGAP- and DFAST-identified pseudogenes that were missed by Pseudofinder were manually inspected in reference to their top BLAST hits: many of these genes appear only marginally shorter than their top homologs (not enough to surpass the 75% length cutoff that we set for the Pseudofinder runs). Additionally, some of these genes did not recruit enough homologs from the reference database to be evaluated as pseudogenes by Pseudofinder. Importantly, both of these criteria can be adjusted by the user in Pseudofinder.

## Conclusions

Overall, we conclude that Pseudofinder is more sensitive towards pseudogene identification than DFAST and PGAP. This sensitivity is due to Pseudofinder including more metrics for pseudogenization (e.g. *dN/dS*); this is particularly apparent in the case of SOPE, whose genome encodes many genes that appear to be under relaxed selection, but have not necessarily acquired obvious inactivating mutations. Nonetheless, the differences identified here, between pseudogene prediction by PGAP, DFAST, and Pseudofinder, demonstrate that identification of pseudogenes is complicated and is best supplemented by manual inspection and parameter optimization. To this end, we put forth Pseudofinder, which offers a standardized pipeline where users can easily tailor parameters relevant to the biological system at hand.

## Methods

### Software Description

Pseudofinder is implemented in Python 3. It has four main built-in commands, or modules: *Annotate*, *Reannotate*, *Sleuth*, and *Visualize*. The *Annotate* command performs the initial pseudogene analysis using a comprehensive database of proteins, such as RefSeq or NR (non-redundant database of proteins), available from NCBI. *Reannotate* is similar to *Annotate*, but allows the user to bypass the most time-intensive step of the pipeline and generate a new set of pseudogene predictions using different parameters. If a closely related reference genome is available, Pseudofinder uses the Sleuth module for single-genome-guided annotation, which can detect relaxed selection (via *dN/dS*), as well as the type and quantity of gene-disrupting mutations in each gene (e.g. frameshift-causing indels, loss of start/stop codons, nonsense mutations, etc.). *Visualize* generates summary plots to assist the user in optimizing parameters for pseudogene identification.

Here we provide descriptions of three of the core modules of Pseudofinder:

- **Annotate**: this module represents Pseudofinder’s core pipeline. It accepts prokaryotic genomes in GenBank format (NCBI compliant, with both gene and CDS features) and a protein sequence database as input, along with many optional parameters. Additionally, users have the option of providing a single reference genome closely related to the query genome, in which case, the *Sleuth* module is invoked. The overall pipeline is outlined in **Figure 2**. First, the input genome is split into coding regions and intergenic regions. Coding regions are predefined in the input annotation, and intergenic regions are defined as the regions between the predicted coding regions. For each coding region, homologs from the reference database are collected using BLASTP (Camacho *et al*., 2009) or DIAMOND (Buchfink *et al*., 2014). Truncated coding regions are identified by comparing gene and alignment lengths to the average lengths of top homologs identified from the reference database. Because genes naturally vary in size, in addition to an arbitrary length cutoff, Pseudofinder will consider the mean and standard deviation of the top DIAMOND/BLAST hits to each queried gene. Fragmented genes are identified as adjacently encoded gene fragments that share the same homologs from the reference protein database (**Figure S1**). For each intergenic region, BLASTX is used to check for significant amino acid sequence similarity in all six reading frames. This process recovers highly degraded pseudogenes that have been missed by gene prediction software, and can also identify regions of pseudogenes upstream or downstream of predicted, truncated gene regions.
- **Sleuth**: while this module is invoked when a reference genome is provided to *Annotate*, *Sleuth* is also a standalone module that accepts as input a prokaryotic genome in FASTA format and a reference genome’s CDS, and performs a pairwise analysis. First, CDS from the reference genome are queried against the genome-of-interest. Homologous regions are then re-aligned using Muscle (Edgar, 2004), and the resulting alignments are processed with respect to indels, nonsense mutations, frameshift-induced early stop codons, loss of start or stop codons, and *dN/dS*. The use of *dN/dS* should be restricted to genomes within a reasonable evolutionary distance (e.g. no more distantly related than at the genus level), and specific genes within a certain evolutionary divergence (e.g. *dS* > 0.01 and *dS* < 3, which are both set as defaults within the software). *Sleuth* also estimates the degree to which frameshift-inducing indels impact the resulting protein sequence: for example, frameshift-causing indels are considered deleterious when they significantly impact the amino acid sequence of the gene product; however, another frameshift shortly downstream can shift the correct codons back into frame, resulting in a negligible impact on the protein sequence. The *Sleuth* module measures the impact that frameshift-inducing indels have on the overall protein sequence and uses this information to predict pseudogenes (**Figure S2**).
- **Visualize**: Annotating pseudogenes requires that we define biologically arbitrary cut-offs. For example, Pseudofinder has a number of parameters that can be tuned by the user, which have the potential to significantly impact pseudogene predictions. These parameters are arbitrary because genes naturally vary in size, number of domains, the amount of frameshift-inducing indels they can tolerate, and their mutation rates. In other words, a one-size-fits-all definition for pseudogenes is not appropriate. We urge users to test multiple settings and visualize each set of results using Pseudofinder’s *Visualize* module (in particular the *dN/dS* and length cutoff compared to reference sequences). This built-in visualization function helps to inform users how their results change as they modify various cut-offs. With a single command, a 3D plot will be generated using Plotly (Plotly Technologies Inc., 2015) to display the number of pseudogenes flagged (z-axis) with any combination of length and similarity parameters (examples available in the following URL: https://github.com/filip-husnik/pseudofinder/wiki/Gallery).

**Figure 2:**
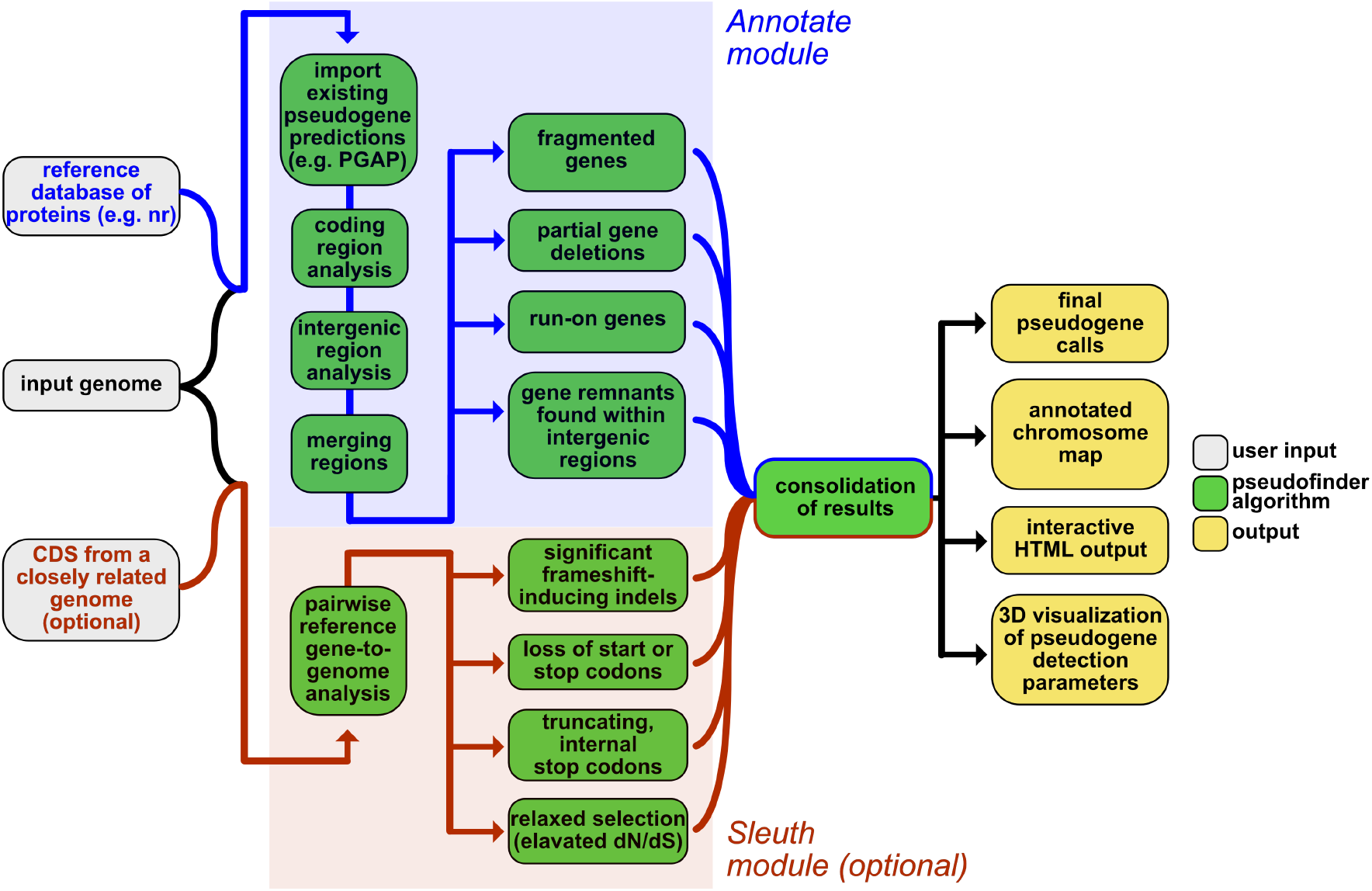
Pseudofinder workflow: the main Annotate branch is shown in the top (blue) part of the workflow, where predicted coding and intergenic regions are compared against proteins from a reference database, allowing the software to identify truncated and run-on ORFs, fragmented genes, and highly degraded gene remnants that lack identifiable gene features. The Sleuth branch is shown in the bottom (red) part of the workflow, where genes from a closely-related reference genome are compared against the genome-of-interest to identify gene inactivations at a finer scale; these inactivations, or gene breakages, can include significant frameshift-inducing indels (i.e. indels that results in substantial changes to the protein sequence), nonsense mutations, loss of start and stop codons, and relaxed selection (elevated dN/dS, measured using PAML [Yang et al., 2007]). Information obtained from these two branches are then consolidated and provided to the user in the form of GFF and FASTA files for downstream processing. Pseudofinder also provides multiple ways for users to visualize the results, including a PDF-formatted genome diagram/map, as well as an HTML-formatted file for interactive exploration of pseudogene predictions.

---

**Supplemental Figure S1:**
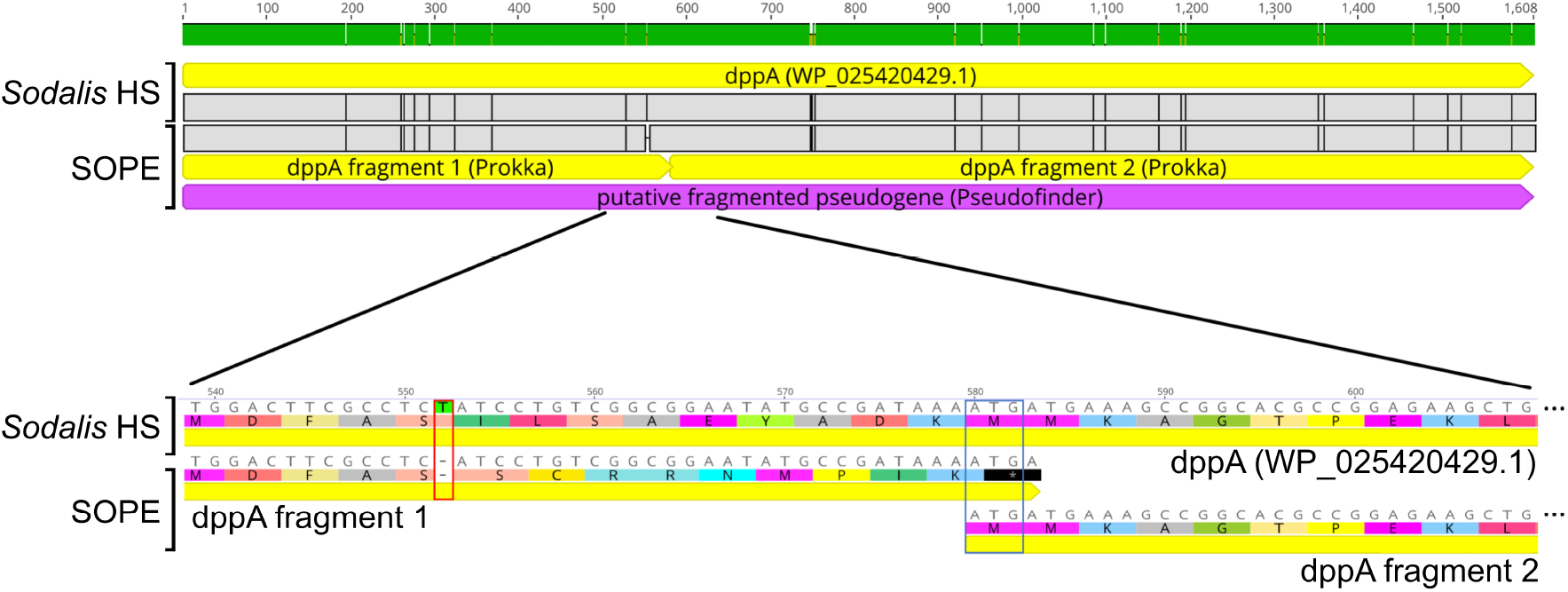
Schematic showing how an alignment looks like between a reference gene and two query genes that have been fragmented. In this example, the dppA gene of SOPE, is compared to its counterpart in the reference genome Sodalis HS. This gene in SOPE was predicted as two different fragments (each with their own open reading frames), both of which align to the N- or C-termini of the reference gene. Pseudofinder will detect these kinds of events and, as part of the output, provide the user with a full, reconstructed gene sequence (shown in purple). The bottom inset shows the nucleotide sequence and corresponding translations, demonstrating how a single-nucleotide deletion (red rectangle at position 552) shifts the frame, resulting in an early, truncating stop codon in position 581, followed by the second predicted fragment of the dppA gene.

**Supplemental Figure S2:**
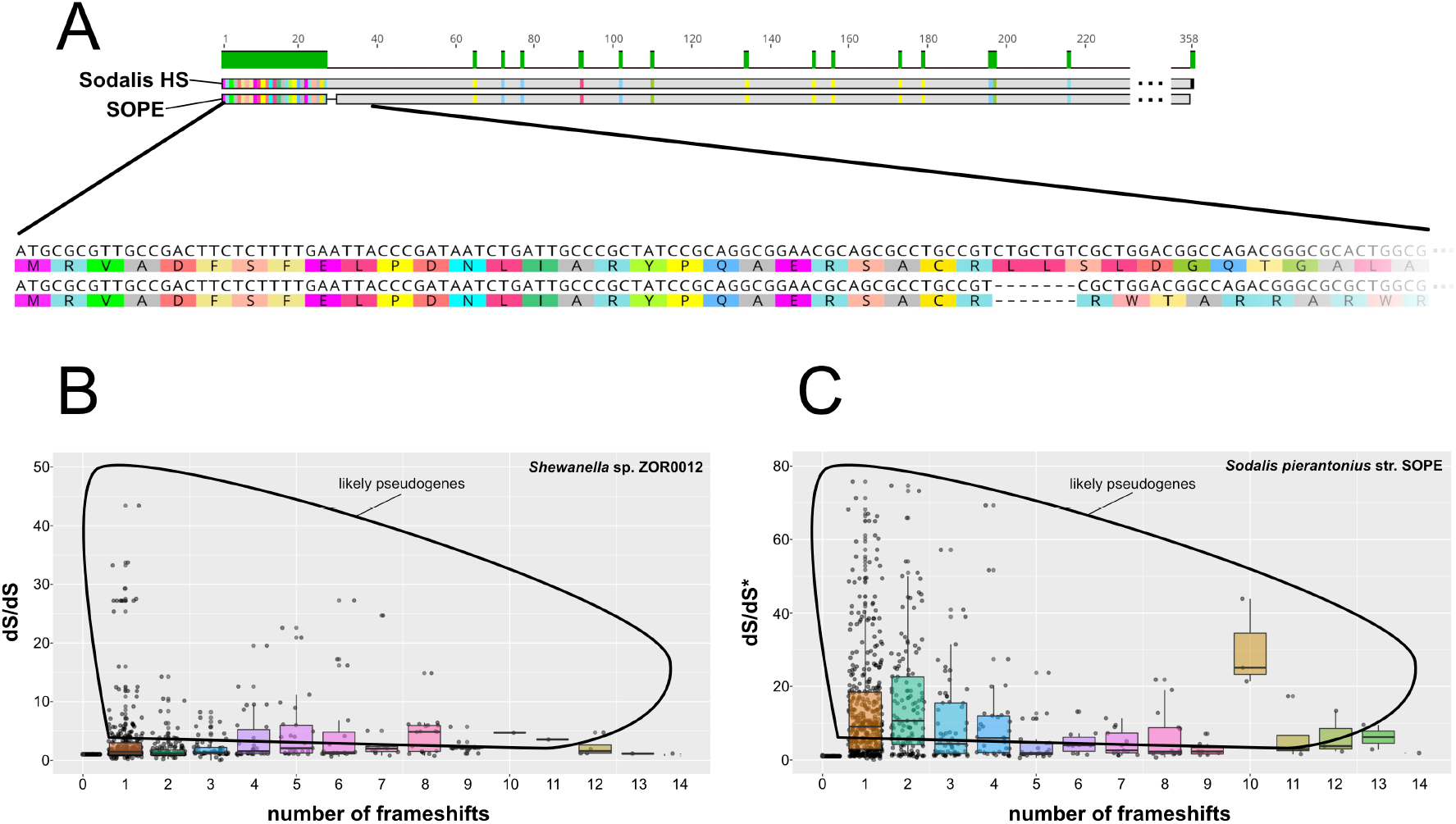
A) Schematic demonstrating how a 7-bp deletion in SOPE’s queA gene (bottom) results in an incorrect translation and a poor downstream alignment. The inset below shows the corresponding nucleotide and peptide alignments. Pseudofinder’s Sleuth module measures the dS value twice: once directly from the nucleotide alignment generated with Muscle (Edgar, 2004), and once from the peptide alignment generated with pal2nal (Suyama et al., 2006) (dS represents the rate of synonymous substitutions per synonymous site, within each gene). The difference between these two dS values is represented in the dS/dS metric, which measures the extent to which frameshift-inducing indels impact the final protein sequence. Panels B and C show the number of frameshift-inducing indels (per-gene), plotted against the impact (dS/dS) of those frameshifts on the overall reading frame of the protein, in B) Shewanella sp. ZOR0012 and C) Ca. Sodalis pierantonius str. SOPE.

